# Oscillatory default mode network coupling in concussion

**DOI:** 10.1101/140368

**Authors:** B.T Dunkley, K. Urban, L. Da Costa, S. Wong, E.W. Pang, M.J. Taylor

## Abstract

**Background:** Concussion is a common form of mild traumatic brain injury (mTBI). Despite the descriptor ‘mild’, a single injury can leave long-lasting and sustained alterations to brain function, including changes to localised activity and large-scale interregional communication. Cognitive complaints are thought to arise from such functional deficits. We investigated the impact of injury on neurophysiological and functionally-specialised resting networks, known as intrinsic connectivity networks (ICNs), using MEG.

**Methods:** We assessed neurophysiological connectivity in 40 males, 20 with concussion, 20 without, using MEG. Regions-of-interest that comprise nodes of ICNs were defined, and their time courses derived using a beamformer approach. Pairwise fluctuations and covariations in band-limited amplitude envelopes were computed reflecting measures of functional connectivity. Intra-network connectivity was compared between groups using permutation testing, and correlated with symptoms.

**Results:** We observed *increased* resting spectral connectivity in the default mode and motor networks in our concussion group when compared with controls, across alpha through gamma ranges. Moreover, these differences were not explained by power spectrum density (absolute changes in the spectral profiles within the ICNs). Furthermore, this increased coupling was significantly associated with symptoms in the DMN and MOT networks – but once accounting for comorbid symptoms (including, depression, anxiety, and ADHD) only the DMN continued to be associated with symptoms.

**Conclusion:** The DMN network plays a critical role in shifting between cognitive tasks. These data suggest even a single concussion can perturb the intrinsic coupling of functionally-specialised networks in the brain and may explain persistent and wide-ranging symptomatology.

## Introduction

Concussion is a common form of mild traumatic brain injury (mTBI), and whilst symptoms are often acute and the majority of patients make a full recovery, a single injury can leave long-lasting and sustained alterations to brain structure and function, even in the absence of symptomatology. A single concussive event has been shown to perturb neurophysiological function, including localised activity and large-scale interregional interactions, with cognitive sequelae manifesting from these phenomenon. Resting functional connectivity is known to be affected, with observed changes in inter-areal spectral correlations (a mechanism that subserves dynamic brain networks for cognition) across multiple frequency scales revealed with magnetoencephalography (MEG). However, the impact of injury on the coordination of neural activity by oscillations in spontaneous and defined *intrinsic connectivity networks’* (ICNs), such as the default mode network, vision, attention, and motor networks, have received little consideration. These innate networks are critical in coordinating and routing information in the brain, and serve functionally-defined and specialised purposes, such as visual perception (Doesburg, Ribary, Herdman, Miller, et al., 2011; Doesburg, Ribary, Herdman, Moiseev, et al., 2011) and directed attention (Jensen, Kaiser, & Lachaux, 2007), amongst others.

Characterising changes in spectral ICNs after concussion could help our understanding of the functional phenotypes of injury and improve diagnostics. Early imaging studies examined alterations to functionally-segregated brain regions and the contribution of these areas to symptoms and specific cognitive deficits (Chen, Johnston, Collie, McCrory, & Ptito, 2007). However, concussion is beginning to be considered an example of disturbed neuronal network communication (Mayer, Mannell, Ling, Gasparovic, & Yeo, 2011). Specifically, there has been a gradual paradigm shift towards thinking of the injury as one of affecting optimal interregional *integration and segregation* – in other words, the coordinated action of areas critical in cognition. The compromised communication within and between networks could lead to a better understanding of the symptomology of concussion. Recent research supports this view and connectivity studies have shown that atypical synchronous network interactions hold power in describing the disorder, in both task-free resting-state, and task-dependent, cognitive-behavioural paradigms (Dunkley et al., 2015; Huang et al., 2016; Pang, Dunkley, Doesburg, da Costa, & Taylor, 2016; Tarapore et al., 2013; Vakorin et al., 2016). The majority of disturbances to the ongoing and spontaneous spatiotemporal interactions of ICN interplay have been found in concussion using fMRI (Johnson et al., 2012), however this technique is limited to measuring ultra-slow fluctuations in blood flow and their regional dynamics. In contrast to fMRI, MEG has the capability to measure higher frequency *neurophysiological* oscillations and the synchrony of these networks which operate at behaviourally-relevant time-scales required for goal-directed cognition and action (Hari & Salmelin, 2012). Specifically, MEG is sensitive to the electrophysiological interactions of primary currents in the brain and can elucidate their frequency composition and dynamics (Baker et al., 2014; Brookes, Woolrich, et al., 2011; de Pasquale et al., 2012) as MEG maps neuronal activity at the millisecond temporal resolution. Rather than simple epiphenomenology, neuronal oscillations are thought to gate information and coordinate the functional coupling of brain areas (Engel, Gerloff, Hilgetag, & Nolte, 2013; Fries, 2005; Stitt et al., 2015; Zumer, Scheeringa, Schoffelen, Norris, & Jensen, 2014). Importantly, perturbations to cortical oscillations and synchronisation are aberrant in a variety of neuropsychiatric disorders, and characterising them provides new understanding of neuropsychopathology.

In concussion studies, MEG has revealed abnormal local changes in neuronal function, including source amplitude (da Costa et al., 2014) and atypical interactions among brain areas (Pang et al., 2016), while further experiments have shown that pattern classification can differentiate those with the disorder from control groups (Huang et al., 2014; Vakorin et al., 2016). In addition, a recent study of persistent concussion symptom patients showed decreased functional connectivity, mediated via phase locking and synchrony, across multiple frequency bands in the default mode network (Alhourani et al., 2016). This evidence suggests brain oscillations contribute to symptoms of the disorder, and large-scale neuronal oscillatory connectivity may explain emergent cognitive sequelae and comorbidities. In a previous study, we took an atlas-guided region-of-interest approach to characterise these metrics and examined how *individual* regions and connections were atypical in their graph properties; in the present paper, we investigated how atypicalities in specialised ICNs and their connectivity are expressed following injury and related to the presence of symptoms.

Our aim was to determine changes in resting brain connectivity mediated by oscillations in established intrinsic brain networks after a single concussion – we predicted that connectivity would be reduced in concussion across multiple frequency scales, and particularly that of the DMN given recent findings (Alhourani et al., 2016). We explored whether these interactions were associated with primary symptoms and secondary conditions (attention, anxiety and depression). We predicted that those with a concussion would express atypical and reduced ICN oscillatory connectivity, and that neurophysiological network interactions would be negatively associated with the presence of primary and secondary symptoms.

## Methods and Materials

### Participants

Resting-state MEG data were recorded from 26 male participants with a single concussive episode (scanned less than three months post-injury, mean days since injury = 32.20, SD = 17.98, mean age at injury = 31.4 years, SD = 6.87). 6 datasets were excluded in the final analysis due to artefactual data (2 due to dental work, and 4 due to excessive head motion during the scan), for a total of 20 participants. The control group consisted of 24 age- and sex-matched participants (mean age = 27.0 years, SD = 5) without any self-reported history of head injury; for final analysis a subset of 20 participants that were most closely matched to the Concussion group on age and IQ were used.

Participants with concussion were identified at the emergency department of Sunnybrook Health Science Centre (Toronto, Canada). Inclusion criteria were: between 20 and 40 years of age; concussion symptoms present during visit to emergency; the MEG scan within 3 months of injury; if loss of consciousness occurred, then less than 30min; if post-traumatic amnesia occurred, then less than 24 hrs; causes of head injury were clear (e.g. sustaining a force to the head); Glasgow Coma Scale ≥13 (within 24 hrs of injury); no skull fracture; unremarkable CT scan; and no previous incidence of concussion. Every participant in the mTBI group was able to tolerate enclosed space for MR brain imaging; be English speaking; be able to comply with instructions to complete tasks during MEG and MR scans; be able to give informed consent. The control group had no history of TBI (mild, moderate or severe) or neurological disorders. Exclusion criteria for both groups included ferrous metal inside the body that might be classified as MRI contraindications, or items that might interfere with MEG data acquisition; presence of implanted medical devices; seizures or other neurological disorders, or active substance abuse; certain ongoing medications (anticonvulsants, benzodiazepines, and/or GABA antagonists) known to directly or significantly influence electroencephalographic (EEG) findings.

All participants underwent brief cognitive-behavioural testing in addition to the MEG resting-state scan. These assessments included: estimates of IQ from the Wechsler Abbreviated Scale of Intelligence (WASI (Wechsler, 1999)); Conner’s Attention-Deficit Hyperactivity Disorder Test; the Generalized Anxiety Disorder 7 test (GAD-7); Patient Health Questionnaire (PHQ9); and the Sports Concussion Assessment Tool 2 (SCAT2 (McCrory, 2009)).

### Procedure and MEG data acquisition

Resting-state MEG data were collected whilst participants were lying supine, and instructed to rest with eyes open and maintain visual fixation on an X within a circle on a screen 60 cm from the eyes. MEG data were collected inside a magnetically-shielded room on a CTF Omega 151 channel system (CTF Systems, Inc., Coquitlam, Canada) at The Hospital for Sick Children, at 600 Hz for 300 seconds. Throughout the scan, head position was continuously recorded by three fiducial coils placed on the nasion, and left and right pre-auricular points.

After the MEG session, anatomical MRI images were acquired using the 3T MRI scanner (Magnetom Tim Trio, Siemens AG, Erlangen, Germany) in a suite adjacent to the MEG. Structural data were obtained as T1-weighted magnetic resonance images using 3D MPRAGE sequences (repetition time [TR] = 2300 ms; echo time [TE] = 2.9 ms; flip angle [FA] = 9°; Field-of-view [FOV] = 28.8 × 19.2 cm; 256 × 240 matrix; 192 slices; 1 mm isovoxel) on a 12-channel head and neck coil. MEG data were coregistered to the MRI structural images using the reference fiducial coil placements. A single-shell head model was constructed for each individual and brain space was normalized to a standard Montreal Neurological Institute (MNI) brain using SPM2.

### MEG data processing

#### Seed definition and virtual electrode output

MEG data were processed using a mixture of the FieldTrip toolbox (Oostenveld, Fries, Maris, & Schoffelen, 2011) and in-house analysis scripts. Time-series were band-pass filtered offline at 1-150 Hz, a Discrete Fourier Transform notch filter applied at the 60 Hz powerline frequency and 2^nd^ harmonic, and a third-order spatial gradient environmental noise-cancellation was applied to the recording. *A priori* sources (seeds) of interest in cortical and sub-cortical regions were identified using coordinates from de Pasquale et al (2012; Visual attention network - VAN, Dorsal attention network - DAN, Default mode network - DMN, Motor network - MOT and Visual network VIS). Figure 1A shows the node locations, Figure 1B shows the analysis pipeline.

**Figure 1.**
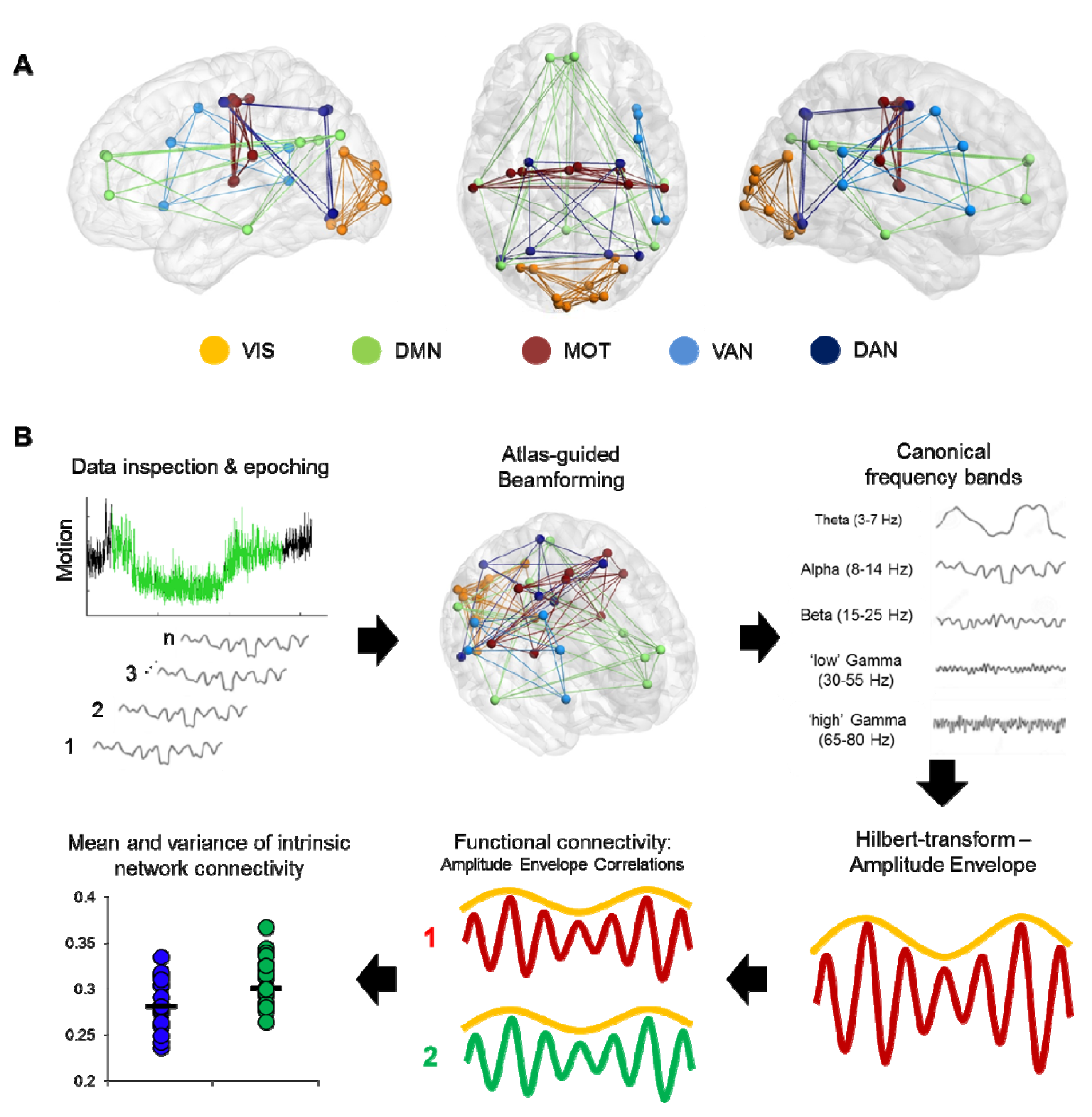
Computing ICN resting-state coupling. **A**. Nodes and internal connections of the predefined ICNs seed regions based on MNI coordinates reported in the literature. **B**. Analysis steps in the pipeline. Data were searched for 2 minutes of minimal head motion from the 5-minute recording, and subsequently epoched into 12 × 10 second segments. A vector beamformer was used to derive ‘virtual-sensor’ time-series from all seeds, and filtered into canonical frequency bands. A Hilbert transform was applied to derive estimates of the instantaneous amplitude envelope. Each pairwise combination of time-series amplitude envelopes inside a network were then correlated to define the degree of connectivity between nodes and averaged per epoch to derive a measure of internal connectivity. These were then averaged over epochs.

Time-series data were reconstructed from these seed locations using a linearly-constrained minimum variance (vector) beamformer (Van Veen, van Drongelen, Yuchtman, & Suzuki, 1997) for each subject and filtered into five canonical bandwidths for further analysis: Theta (4-7 Hz), Alpha (8-14 Hz), Beta (15-30 Hz), Gamma (30-55 Hz) and High-Gamma (65-80 Hz). A beamformer is a spatial filter used to suppress signals from extrinsic neural and noise sources, whilst maintaining unit gain for activity in a target brain location (in this case, the defined seed locations). Individual weight vectors are applied to each sensor measurement and summated to give estimated source activity at target seed locations. This type of spatial filter is also effective at suppressing ocular artefacts generated by eye movements, and non-ocular artefacts, such as cardiac and muscle activity (Cheyne, Bostan, Gaetz, & Pang, 2007; Muthukumaraswamy, 2013).

### Statistical analysis

Each of the analyzed frequency ranges from each subject were then submitted to a functional connectivity analysis, by computing amplitude envelope correlations (AEC) across each of the 10 second epochs, based on the instantaneous amplitude estimate of each sample from the filtered time-series calculated using the Hilbert transform. The magnitude of AEC between all pairwise combinations of the seeds varied between 1 (perfect correlation) and −1 (perfect anti-correlation). These values quantify the time-varying correlation in the envelope between any two sources, referred to henceforth as functional connectivity.

Then, adjacency matrices with AEC values acting as edge weights for all sources pairs were constructed, which resulted in a matrix of weighted undirected graphs in each analysed frequency band for each participant. Connectivity weights between seeds within an intrinsic network were averaged to characterise the magnitude of spontaneous intra-network coupling. These were then either averaged over the 12 epochs (2-minute run) to derive time-averaged connectivity, or the standard deviation was calculated to define temporal dynamism (Koelewijn et al., 2015; Muthukumaraswamy, 2013), albeit in network connectivity, rather than local source oscillatory amplitude. These adjacency matrices were then divided into the respective groups and inferential statistics investigating group differences for mean edge weight were implemented using non-parametric permutation testing (20,000 iterations), which do not require the data distributions to be normal. False positives due to multiple comparisons were controlled using Bonferroni-correction across frequency-bands. Cognitive-behavioural correlation analyses were conducted using the MATLAB Statistics Toolbox (The Mathworks, Inc.). Networks were visualized and figures produced using BrainNet Viewer (Xia, Wang, & He, 2013).

## Results

### Comorbid symptoms in concussion

All clinical outcome measures were greater in the Concussion group than the Control group (Conners, t = 2.52, p = 0.02; GAD7, t = 2.44, p = 0.02; PHQ9, t = 3.36, p = 0.002; SCAT2 symptoms, t = 4.53, p = <0.001; SCAT2 severity, t = 3.09, p = 0.004), whilst being matched on the WASI (t = −1.79, p = 0.08). Table 1 shows Concussion group demographic and injury information, including symptom number and severity, days since injury, whether loss of consciousness occurred and for how long, Glasgow Coma Scale score, presence of post-traumatic amnesia, and mechanism of injury. Symptom severity was most associated with and explained the greatest degree of variance in anxiety (R^2^ = 0.60, p < 0.001), followed by depression (R^2^ = 0.28, p < 0.001), and finally attention symptoms (R^2^ = 0.27, p < 0.001).

**Table 1.**
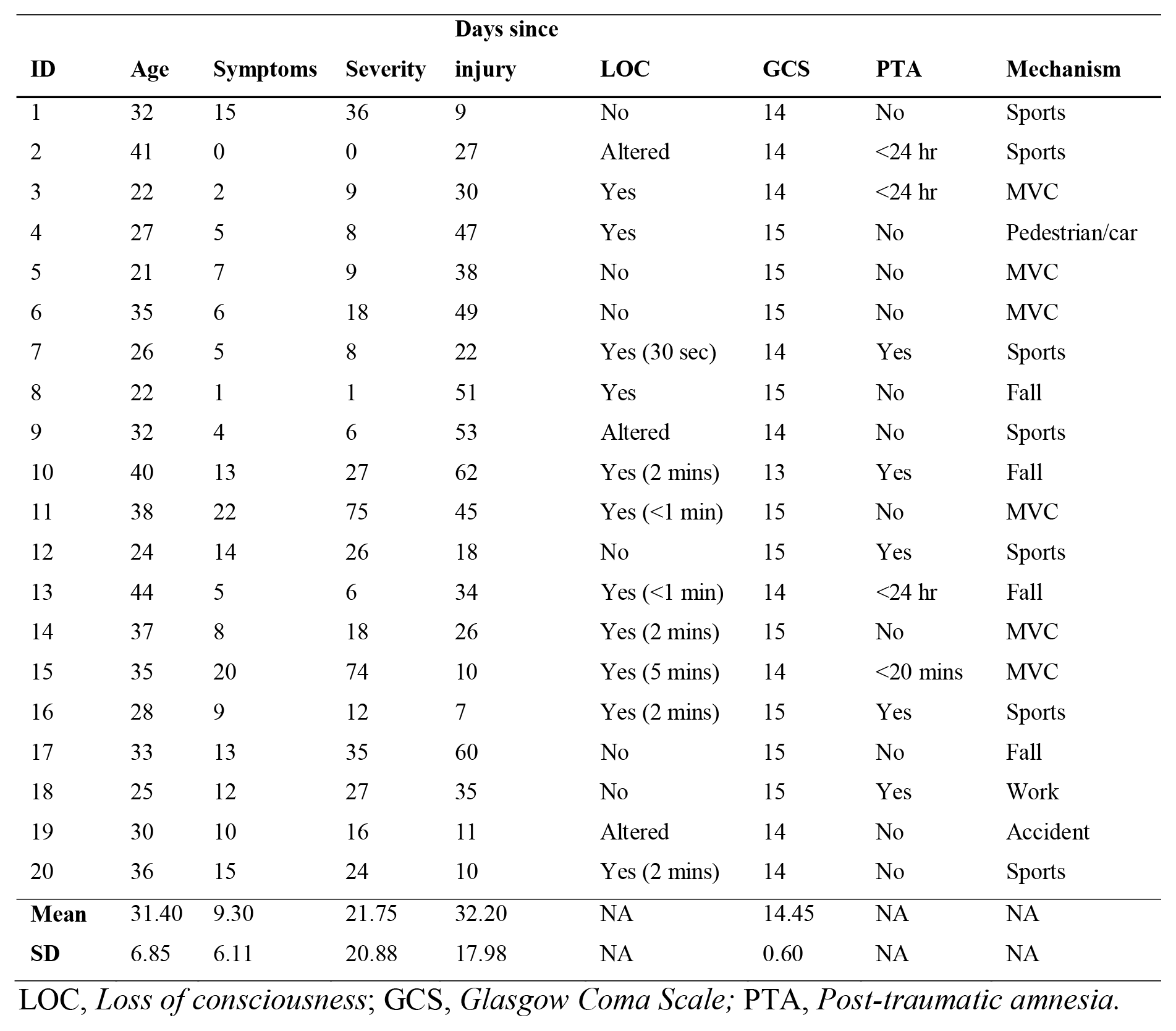

### Increased DMN and MOT network resting connectivity following Concussion

Significant increased DMN connectivity was observed in the alpha and beta ranges for the Concussion group compared to Controls (Bonferroni-corrected within frequency bands at p < 0.05; null distributions generated using 20,000 permutations, Figure 2). Connections between the ventro-medial prefrontal cortex (vmPFC), and the dorso-medial prefrontal cortex (dmPFC) and right medial prefrontal cortex (RMPFC) appear to drive this difference, exhibiting the greatest degree of hypercoupling in the Concussion group compared to Controls. Elevated coupling in Concussion was also found in the MOT network across the alpha, beta and gamma ranges (p_corrected_ < 0.05). No significant differences were observed in the standard deviation of internal coupling across epochs (all p’s > 0.05).

**Figure 2.**
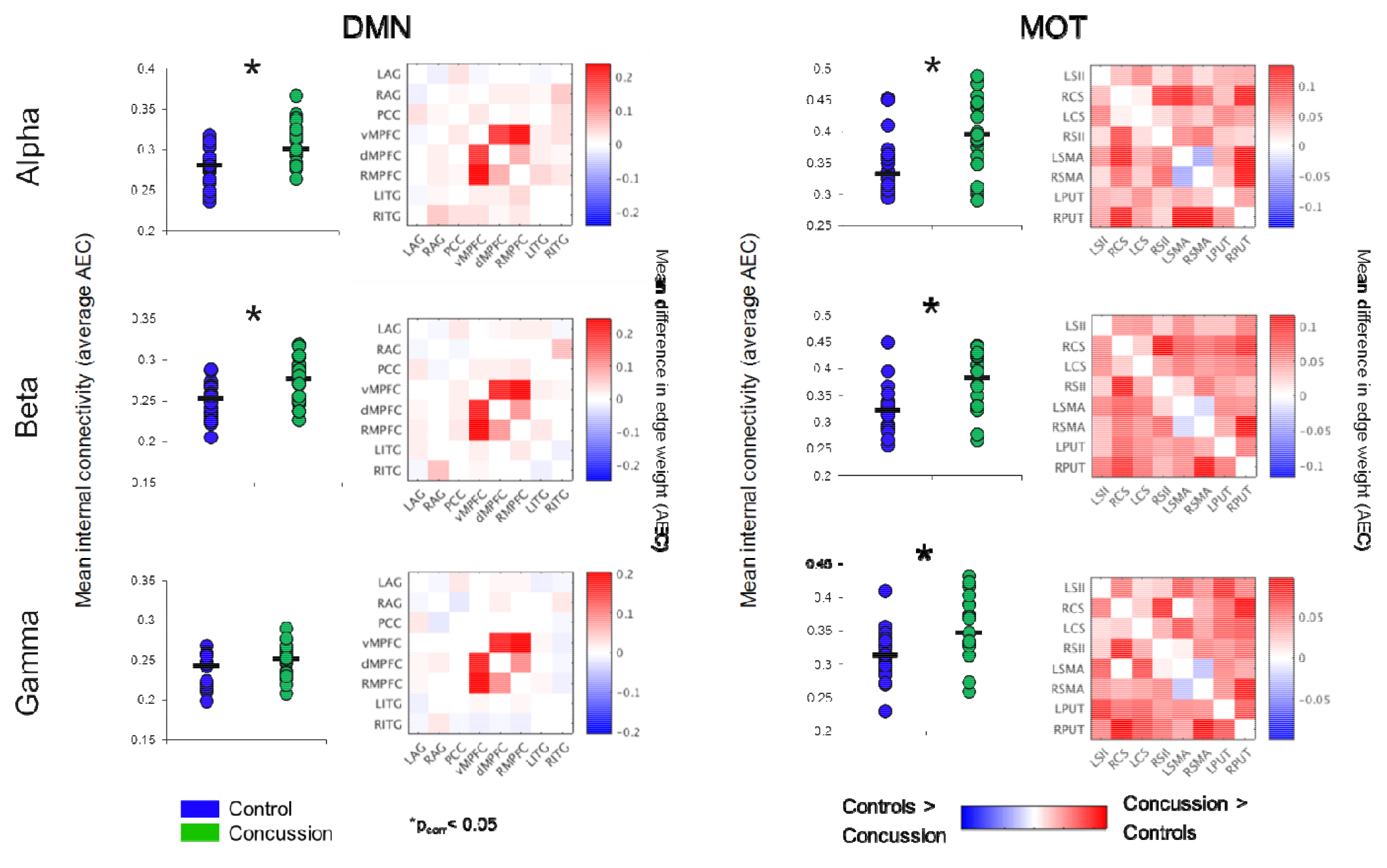
Band-specific ICN connectivity mediated by amplitude envelope correlations. Scatter plot dots show intra-subject mean across epochs, black line represents median over participants for each group in the DMN (left column) and MOT (right) networks, in the alpha (top row), beta (middle row) and gamma ranges (bottom row). *p_corrected_ < 0.05. Connectivity matrices show edge weight differences (Concussion minus Controls) in respective ICNs – contrasts revealed the DMN increases in the Concussion group were driven by VMPFC to RMPFC and dMPFC connections.

### Resting-state spectral power does not explain differences in ICN coupling

To determine the extent to which changes in intra-network coupling were dependent on oscillatory power, the mean internal power spectrum for each of ICNs was calculated and divided into canonical frequency ranges – qualitative assessment of the spectrum show an apparent increase in low-frequency power/an alpha peak shift towards low-frequencies, particularly within the DMN (Figure 3). Mixed ANOVAs on each of the ICNs independently revealed a main effect of band (p < 0.001), as expected, but not of group or any interaction (p > 0.05). Post-hoc contrasts between groups within bands revealed no significant differences (all p’s > 0.05). This suggests connectivity in the bands exhibiting between group differences is not explained by raw spectral power.

**Figure 3.**
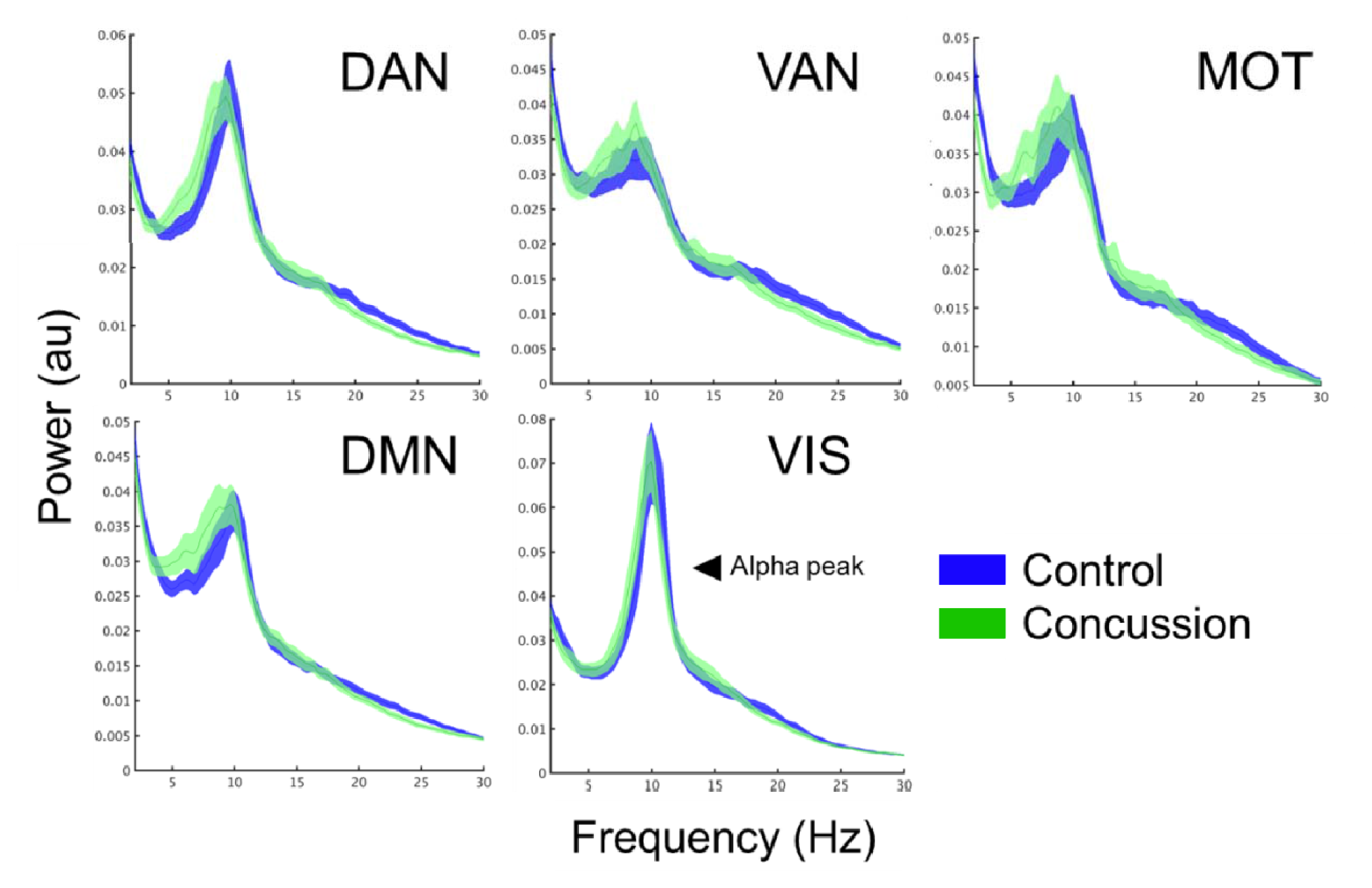
ICN spectral power content, averaged over node regions that comprise a network. Mean whole-network spectral power content with ±1 standard error bars for the Concussion (green) and Control groups (blue). Of note is the 10 Hz alpha peak, prominent in the visual network.

### DMN connectivity correlates with symptom number

In addition to the main MEG findings, we also conducted follow-up analyses of the relation between network connectivity and concussion symptom number (Figure 4 shows scatter plots of connectivity versus symptom presence, and test-statistics for full and partial correlations are detailed in Table 2). Specifically, we examined brain-behaviour relations in the DMN and MOT networks across frequency ranges where differences were observed in between groups contrasts; the alpha, beta and gamma band (Bonferroni-corrected across frequencies). Significant correlations (non-parametric Spearman’s Rho) with symptoms were observed in the DMN and MOT across alpha, beta and low gamma ranges; however, partial correlations with comorbidity symptom scores (Conner’s, GAD7, and PHQ9) entered covariates revealed that the variance in MOT connectivity was not solely driven by concussion symptoms. Factoring in these covariates reduced the full correlation coefficients, such that they were found to no longer be significant – critically, the DMN correlations remained significant at the alpha and beta frequencies (Figure 4 shows original scatter plots linear least squares regression line).

**Figure 4.**
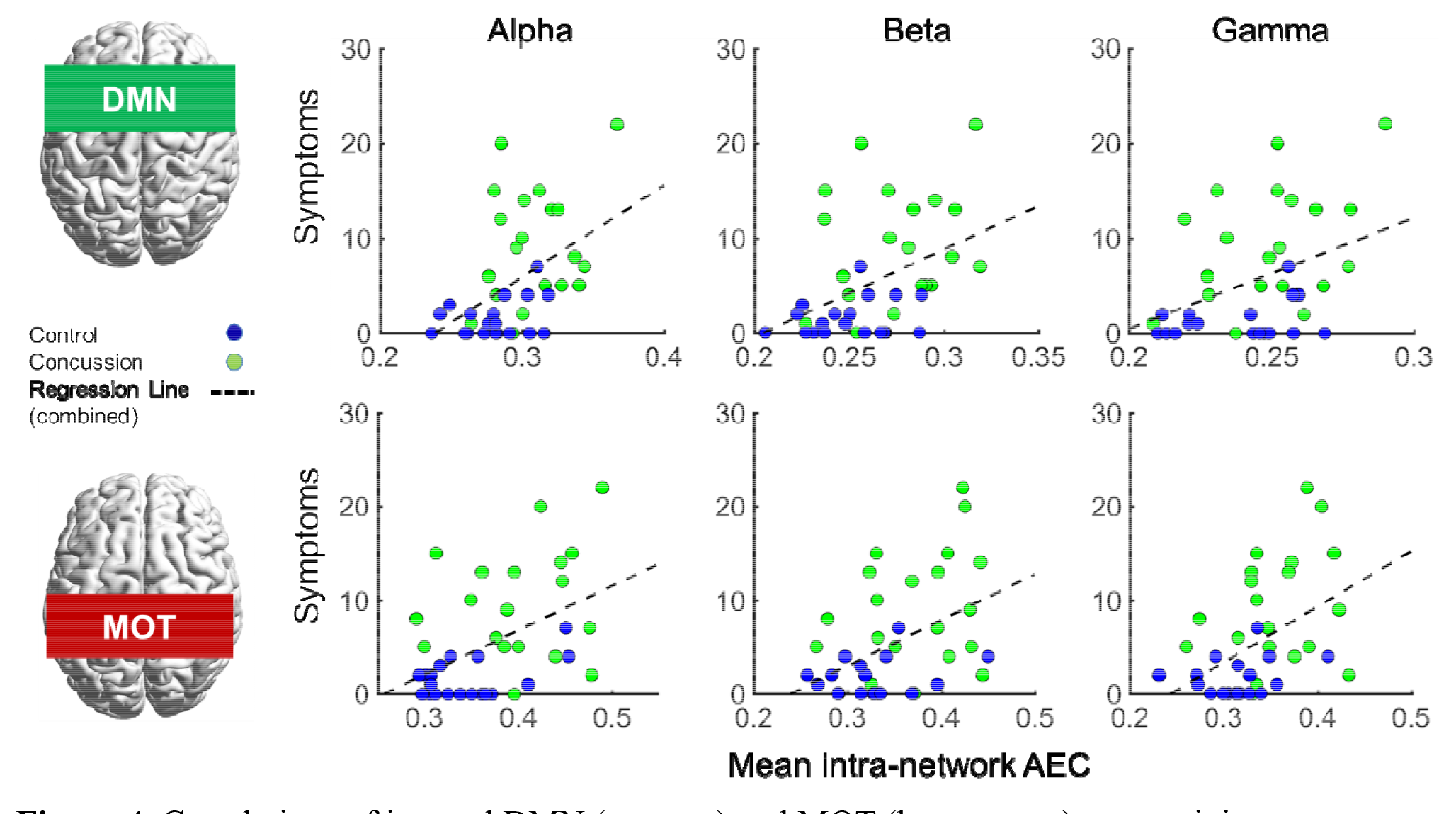
Correlations of internal DMN (top row) and MOT (bottom row) connectivity versus Concussion symptoms in the alpha, beta and gamma ranges. Scatterplots show original (full) correlations and least-squares fit line; test statistics for full and partial correlations are given in Table 2.

**Table 2.**
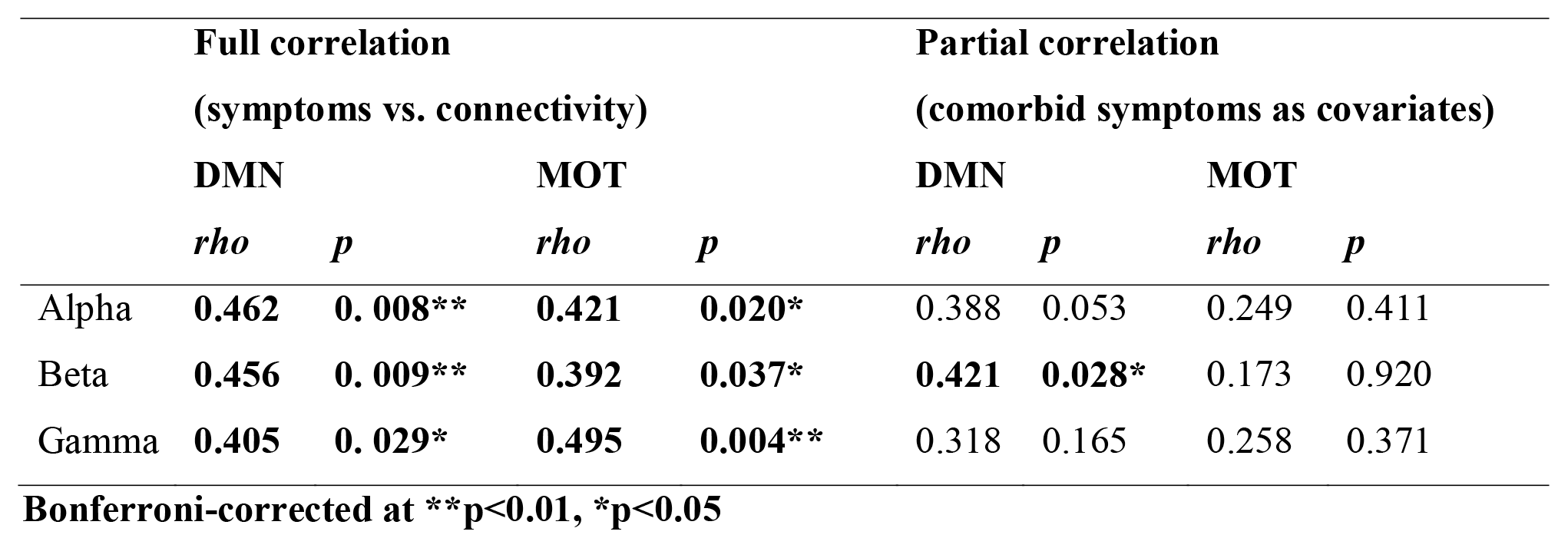

## Discussion

### Summary

In this study, we used MEG to investigate frequency-specific interactions within functional ICNs in adults with a single concussion compared to a matched control group. In contradiction to our initial predictions, a single concussion was associated with increased functional connectivity mediated via band-limited AEC – specifically, elevated coupling within DMN and motor networks, and importantly, in the absence of canonical band power spectrum differences. Intra-DMN connectivity was positively associated with concussion symptoms, even when controlling for secondary/comorbid outcomes (depression, anxiety and attentional problems), which suggests that the internal coupling of this task-negative (Raichle et al., 2001) and dynamic ‘cortical-core’ network (de Pasquale et al., 2012) is particularly prone to the effects of even relatively mild traumatic brain injuries, which are to a degree independent of comorbidities that follow. Additionally, elevated MOT connectivity was also associated with symptoms and secondary sequelae.

### Elevated Default Mode Coupling in Concussion

We found altered oscillatory-mediated connectivity in the DMN in concussion patients. These results appear in opposition to previous fMRI literature examining BOLD fluctuations (Johnson et al., 2012; Mayer et al., 2011; McDonald, Saykin, & McAllister, 2012; Zhou et al., 2012). The DMN, known as the primary resting state network, involves coordinated action between the posterior cingulate cortex (PCC), bilateral angular gyrus, ventral medial prefrontal cortex (vmPFC), dorsal medial PFC (dmPFC) and inferior temporal gyrus. Studies evaluating the slow frequency haemodynamic components have shown overall reductions in connectivity of the DMN in mTBI, especially in posterior regions such as in the PCC, inferior parietal and precuneus (Kou & Iraji, 2014; Mayer et al., 2011). In contrast, we found overall *increased* mean coupling within the default mode. Further evaluation of internal edge weights found that these changes were mainly driven by hyper-coupling between the vmPFC, dmPFC and rmPFC frontal regions, in contrast to the posterior bias reported in fMRI work.

This study shows the importance of using MEG which neurophysiological mechanisms underlying resting state networks. While it is thought that the ultra-slow BOLD is recapitulated and positively correlated with AEC in alpha and beta bands in MEG (Brookes et al., 2014; Brookes, Hale, et al., 2011; Brookes, Woolrich, et al., 2011), this is possibly not true in brain injury where the mechanisms of interregional coupling (that is phase based or envelope coupled) do not function as expected. Apparently conflicting observations - that DMN connectivity in fMRI and MEG positively correlate in healthy participants, but in concussion, patients show differential connectivity across modalities, *decreased* in fMRI and *increased* in MEG – could be due to the impact of injury on distinct mechanisms, such as haemodynamics and neurovascular coupling, changes in regional or global cerebral perfusion, disruption to the neurometabolic balance, and oxidative stress (i.e. endogenous changes in neuronal excitability). Other recent MEG findings show that chronic mTBI patients exhibit *decreased connectivity* in phase-synchronised DMN measures (Alhourani et al., 2016), which further suggests concussion affects neurophysiological fingerprints. These varying results should be considered in light of the fact that all these measures of connectivity differ, (amplitude envelope coupling vs. phase synchrony & BOLD correlations), and capture distinct neural mechanisms in brain communication (Engel et al., 2013; Siegel, Donner, & Engel, 2012). Moreover, the time since injury to scan was different in all studies, capturing either acute/subacute (as here), or chronic stages (Alhourani et al., 2016). These different results could perhaps be explained by compensation mechanisms – that during the acute/subacute recovery, the hyperconnectivity is a reflection of an attempt to reorganise networks. Clearly, these results suggest a multimodal imaging and a synergistic approach to connectivity is required to explain the complexity of pathology in concussion.

The DMN is a task-negative network that contains ‘hubs’ that facilitate task-switching and engagement of other functional networks (de Pasquale et al., 2012), such as those linked to memory, attention and executive function (Buckner, Andrews-Hanna, & Schacter, 2008; Mason et al., 2007). These are cognitive domains oft-reported to be dysfunctional in concussion (Baillargeon, Lassonde, Leclerc, & Ellemberg, 2012; Tapper, Gonzalez, Roy, & Niechwiej-Szwedo, 2016; van der Naalt, van Zomeren, Sluiter, & Minderhoud, 1999), and in light of the DMN’s role in cognition and network toggling, the hyperconnectivity we report might reflect an inability to disengage the DMN in response to shifting task demands, and explain some of the emergent cognitive deficits seen here. The DMNs critical role in neuropsychopathology and cognitive deficits is further supported by studies of other disorders, including ADHD (Lansbergen, Arns, van Dongen-Boomsma Martine, Spronk, & Buitelaar, 2011), Alzheimer’s disease (Stam et al., 2006, 2009), PTSD (Dunkley et al., Huang et al.), ASD (Ye et al., 2014; Yerys et al., 2015), depression (Nugent, Robinson, Coppola, Furey, & Zarate, 2015), and schizophrenia (Camchong, MacDonald, Bell, Mueller, & Lim, 2011).

Oscillations underlie communication processes between distinct brain regions, (i.e. within or between brain network connectivity) (Engel et al., 2013; Fries, 2005, 2015). The complex interplay among regions that comprise the DMN, and ICNs more generally, and how they are mediated by frequency-specific connectivity mechanisms, further highlights the important role of oscillations and their contribution to network architecture and interregional communication – perhaps most importantly, these results show that they are susceptible to perturbations through injury. These neurophysiological processes are intrinsically linked to the distinct topological patterns of communication that constitute functional resting networks (Brookes, Woolrich, et al., 2011; Hunt et al., 2016). Any disruption in the internal balance of these networks may impact the ability for them to switch, engage, or separate - altering cognitive efficiency or recruitment processes. For example, the task-positive executive network is active during externally directed behaviour and requires communication between bilateral DLPFC and PCC. However, in the concussion cohort the connectivity patterns of these brain regions was altered and thus we speculate that task related activity would also be effected. We believe these observations show dysfunctional elevated integration within networks, and an inability to efficiently segregate/decouple areas (Dunkley et al., 2015). Thus, a disruption in the internal balance could lead to altered cognition, which is consistently shown to occur following concussion (Banks et al., 2016; Brown, Dalecki, Hughes, Macpherson, & Sergio, 2015; Tapper et al., 2016).

### Increased Motor Network Connectivity

In addition to our DMN observations, we also found that the MOT network showed multiscale network dysfunction, including the beta band. Beta oscillations have been shown to serve a critical role in long-range synchrony (Engel & Fries, 2010), with the connectivity of these ICNs related to structural connections linking distinct anatomical areas (Park & Friston, 2013); these connections rely on white matter tracts to facilitate signal conduction and increase velocity (Voets et al., 2012). In patients with Parkinson’s disease, increases in resting-state MOT beta connectivity has been related to slower reaction times during visual-motor related tasks (Pfurtscheller, Zalaudek, & Neuper, 1998) and similar changes to reaction time are evident in concussion (Warden et al., 2001). However, the increased MOT connectivity could be a general indicator of functional changes following concussion (e.g. fatigue), and not specific to the symptom scale we used in this study.

Increased beta in the MOT may be due to a variety of neural circuitry changes, possibly related to GABA concentration (Gaetz, Edgar, Wang, & Roberts, 2011). Changes to spectral connectivity could be the result of myriad neurophysiological changes, including altered inhibitory input and changes to excitatory threshold which could alter the connectivity and oscillatory patterns among the brain regions within the MOT. In the motor cortex, beta oscillatory power is high during rest and decreases during movement, known as the event-related-beta desynchronization (ERBD), related to local processing during movement (Pfurtscheller et al., 1998). In Parkinson’s, the increase in beta oscillations in MOT are related to dampened neural activation during motor activation, and slower reaction times during tasks.

As neural oscillations are thought to be a managed through a relationship between excitatory and inhibitory neurotransmission and related to resting GABA concentration (Gaetz et al., 2011) the increased beta amplitude coupling within the MOT could be reflective of an increase in cross-regional inhibitory processes, that could be the cause of increased motor threshold (Tallus, Lioumis, Hämäläinen, Kähkönen, & Tenovuo, 2012), and inhibition and excitability in motor cortex, as measured by transcranial magnetic stimulation (TMS) in concussion (Pearce et al., 2014). Furthermore, increased GABA concentration, an inhibitory neurotransmitter in the motor cortex has been reported in animal and human studies of concussion (Guerriero, Giza, & Rotenberg, 2015; Tremblay et al., 2014). Although these changes in circuitry are evident at the neuronal level, they will also impact short and long range connectivity patterns to which MEG is sensitive.

Local alpha oscillations and their regional coupling are thought to facilitate information integration and segregation within and across neural populations via modulation and gating of local inhibition and excitation (Doesburg, Green, McDonald, & Ward, 2009; Jensen & Mazaheri, 2010). These multi-frequency aberrations suggest disruption to communication channels, and align with mechanism of injury where high tensile and sheering forces have been shown to alter white matter connections and therefore local neural activity and large-scale synchrony. Thus, the functional deficits from injury may alter the coordination of information transfer and integration across a variety of complex processes, including oscillatory coupling (investigated here), and phase-amplitude coupling and cross-frequency interactions to which MEG is sensitive (Baillet, 2017; Florin & Baillet, 2015) and have shown to be affected by concussion (Antonakakis et al., 2016). As such, the heterogeneous nature of concussion can lead to various changes in brain function on local and global scales, which aligns with the results we found of spectral connectivity changes and reinforces the fact that a multimodal, multi-analytical approach is required to comprehensively describe neurophysiological and anatomical changes after a concussion.

### Relations between functional connectivity and structural architecture

Cartography of the human ‘connectome’ has seen a surge in recent years, but progress has been slow in elucidating the association between electrophysiological connectivity, and structural connectivity, as in cortical oligodendrocytes and myelin content. Preclinical work using optogenetics to stimulate neuronal (electrical) activity has shown that this promotes adaptive myelination (Gibson et al., 2014), and a recent human study has proposed that there is a strong link between neural connectonomics, their dynamics and cortical white matter structure (or ‘myeloarchitecture’ (Hunt et al., 2016)). It seems reasonable to posit, given these findings, that our observations in altered connectivity of cortical ICNS could be related to neuromorphological changes (Sussman et al., 2017), and more specially microscopic white matter pathology following concussion, including demyelination and/or differentiation (Huang et al., 2012; McAllister et al., 2012), reductions in cortical thickness (Hamberger, Viano, Säljö, & Bolouri, 2009; Lewén et al., 1999; Urban et al., 2016), and Wallerian degeneration (Mckee & Daneshvar, 2015; Smith, Hicks, & Povlishock, 2013).

Several hypotheses have been proposed to better understand the underlying neurophysiological causes of increases in resting state connectivity. One frame of thought suggests that connectivity increases could be a result of compensation for injury. Specifically, hyper-connectivity could be representative of recruitment of additional resources to maintain cognitive function – the elevated connectivity in frontal regions of the DMN in the concussion group aligns to the general findings of frontal susceptibility to injury (for a review, see (Eierud et al., 2014)). The disruption to white matter tracts could cause changes in the long range connections (such as between PCC and frontal regions) and result in increases in small-world connectivity. This can be seen in our results across the DMN, particularly that increases in spectral connectivity can be attributed to the hypercoupling in frontal regions.

### Conclusion

A particular focus in concussion research is to better understand the link between symptom severity/presence and altered brain function, both in the short and long term. We found a significant relation between symptoms and increases in spectrum connectivity in the DMN and MOT. The association between DMN and symptoms was maintained even while controlling for secondary factors as anxiety and depression. Given the critical role of the DMN in cognitive functions, these abnormal patterns may well underlie the cognitive difficulties so often reported following concussion. Finally, these results, when combined with observations from other studies, suggest MEG may be able to identify electrophysiological changes along the temporal continuum of recovery, from the short term, acute stages of injury, through to later, chronic phases. In conclusion, longitudinal MEG studies of neurophysiological function may be able to link symptoms with intrinsic function and predict the course of recovery in individuals.

## Acknowledgements

Thanks are given to Amanda Robertson, Allison Bethune and Marc Lalancette for help in the data collection, and Simeon Wong for help with the data pre-processing. This work was supported by funding from Defence Research and Development Canada (DRDC) (Contract #: W7719-135182/001/TOR) and the Canadian Forces Health Services to MJT and EWP.

## Financial disclosures and competing interests

The authors declare no competing financial interests or potential conflicts of interest.

